# Colorectal cancer heterogeneity co-evolves with tumor architecture to determine disease outcome

**DOI:** 10.1101/2025.08.11.669722

**Authors:** Fabien Bertillot, Solène Hervé, Nelly Drobjazko, Noemi Kedei, Mona El Gendi, Maria O. Hernandez, Kai Kruse, Erdogan Pekcan Erkan, Mirjam Binner, Karolina Punovuori, Teijo Pellinen, Maarit Ahtiainen, Jan Böhm, Jukka-Pekka Mecklin, Caj Haglund, Jaana Hagström, iCAN, Mike Nikolaev, Nikolce Gjorevski, Matthias Lütolf, Toni T. Seppälä, Sara A. Wickström, Yekaterina A. Miroshnikova

**Affiliations:** Department of Cell and Tissue Dynamics, Max Planck Institute for Molecular Biomedicine, 48149 Münster, Germany; Stem Cells and Metabolism Research Program, Faculty of Medicine, University of Helsinki, 00290 Helsinki Finland; Laboratory of Molecular Biology, National Institute of Diabetes and Digestive and Kidney Diseases, National Institutes of Health, Bethesda, Maryland 20892, USA; Max Planck Research Group Biophysical Regulation of Cell State Dynamics, Max Planck Institute for Molecular Biomedicine, 48149 Münster, Germany; Spatial Imaging Technology Resource Office of Science and Technology, Center for Cancer Research, National Cancer Institute, Bethesda, MD 20892, USA; Max Planck Institute for Molecular Biomedicine, 48149 Münster, Germany; Department NEON-S, Faculty of Medicine and Health Technology, University of Tampere, Tampere, Finland; Institute for Molecular Medicine Finland, Helsinki Institute of Life Science, University of Helsinki, Helsinki, Finland; Central Finland Biobank, Hospital Nova of Central Finland, Well Being Services County of Central Finland, Jyväskylä, Finland; Department of Pathology, Hospital Nova of Central Finland, Well Being Services County of Central Finland, Jyväskylä, Finland; Faculty of Sport and Health Sciences, University of Jyväskylä, Jyväskylä, Finland; Department of Education & Research, The Wellbeing Services of Central Finland, Jyväskylä, Finland; Research Programs Unit, Translational Cancer Biology Program, University of Helsinki, Helsinki, Finland; Department of Surgery, University of Helsinki and Helsinki University Hospital, Helsinki, Finland; Department of Oral Pathology and Radiology, University of Turku, Turku University Hospital, Turku, Finland; Department of Pathology, University of Helsinki, FI-00014 Helsinki, Finland; iCAN Digital Precision Cancer Medicine Flagship, University of Helsinki; Institute of Human Biology (IHB), Roche Pharma Research and Early Development, Roche Innovation Center Basel, Basel 4052, Switzerland; Applied Tumor Genomics, Research Programs Unit, University of Helsinki; Department of Gastroenterology and Alimentary Tract Surgery, Tampere University and Tays Cancer Centre, Tampere University Hospital; Department of Gastrointestinal Surgery, Helsinki University Central Hospital, University of Helsinki, Helsinki, Finland; Helsinki Institute of Life Science, Biomedicum Helsinki, University of Helsinki, FI-00290 Helsinki, Finland

## Abstract

Intratumoral heterogeneity, originating from genetic, epigenetic, and phenotypic cellular diversity, is pervasive in cancer. As these heterogeneous states employ diverse mechanisms to promote tumor progression, metastasis, and therapy resistance, emergence of cancer heterogeneity is one of the most significant barriers to curative treatment. Here we leverage deep learning approaches to develop a high-throughput image-analysis paradigm with subcellular resolution that quantifies and predicts colorectal cancer (CRC) patient outcome based on tissue architecture and nuclear morphology. We further combine this approach with spatial transcriptomics, multiplex immunohistochemistry, and patient-derived organoids to uncover a dynamic, co-evolutionary relationship between tumor architecture and cell states. We identify clinically relevant architectural interfaces in CRC tissue that diversify cellular identities by favoring distinct cancer stem cell states and thus promote evolution of tumor heterogeneity. Specifically, tissue fragmentation and associated compressive forces promote loss of classic stem cell signature and acquisition of fetal/regenerative stem cell states, which initially emerges as a hybrid state with features of epithelial-to-mesenchyme (EMT) transition. Additional tumor stroma communication then diversifies these states into distinct stem cell and EMT states at the invasive margins of tumors. Reciprocally, Wnt signaling state of tumor cells tunes their responsiveness to tissue architecture. Collectively, this work uncovers a feedback loop between cell states and tissue architecture that drives cancer heterogeneity and cell state diversification. Machine learning-based analyses harness this co-dependency, independently of mutational status or cancer stage, to predict patient outcome.

## Introduction

Heterogeneity is a highly prevalent feature of human cancers. Most solid tumors are complex ecosystems of genetically, epigenetically and phenotypically heterogenous cell states/types, surrounded by morphologically and compositionally diverse cellular and acellular stroma. Importantly, intratumor heterogeneity associates with poor patient prognosis, presumably due to the diversification of cellular states ^1,2^. A substantial proportion of this heterogeneity is independent of secondary mutations within the tumor ^3^. However, whether the cellular state evolution directly leads to the emergence of morphologically distinct tissue organization, or if the tissue microenvironment, due to its ability to impart biochemical and mechanical signals, can drive cell state heterogeneity remain fundamental open questions. Furthermore, how these cell-intrinsic and cell-extrinsic mechanisms interact to promote cancer aggression and co-evolve during tumor development is unclear.

The intestinal epithelium is the fastest self-renewing tissue in mammals and harbors a pool of adult stem cells to facilitate its turnover ^4^. Importantly, these tissue-resident stem cells that play pivotal roles for intestinal homeostasis and regeneration during wounding, are also the cells of origin and drivers of progression in colorectal cancer (CRC). Recent findings indicate that intestinal stem cells in CRC exist in a dynamic continuum between adult stem and fetal-like and regenerative stem cell states that are normally detected during intestinal development or tissue damage ^5^. During CRC progression, adult stem cell signatures become depleted, while primitive fetal or regenerative programs re-emerge and expand upon metastasis or in response to treatment ^5–7^. Such stem cell state plasticity leads to high intra- and inter-tumor heterogeneity and severely hampers targeted strategies. While the precise molecular programs that control these states are complex and heterogeneous, detachment from the main tumor mass and initiation of metastases seems to require (or be induced by) the loss of classic marker of colonic stem cells, the leucine-rich repeat-containing G-protein coupled receptor 5 (LGR5), the activation of TGFβ1 and YAP signaling, and acquisition of EMP1, Annexin A1 (ANXA1), PLAUR and other fetal stem cell state markers ^5–8^. Thus, understanding the molecular mechanisms of these cell state interconversions is critical for understanding the mechanisms that drive CRC progression.

Decades of research have uncovered the key genetic drivers of CRC, including mutations in BRAF, KRAS, and microsatellite instability (MSI), that are also used to clinically define therapeutic strategies ^9^. In addition, emerging evidence suggests that phenotypic cellular plasticity that may arise independently of secondary genetic mutations and could be driven by epigenetic changes and transcriptional plasticity ^10,11^. Besides these genetic and epigenetic alterations, CRC, like most cancers, is associated with changes in the microenvironmental composition, such as immune cell infiltrates and emergence of a cancer associated fibroblast (CAF) populations that contribute to cancer cell behavior and disease progression ^12–14^. In addition to changes in cell composition, the tumor and its microenvironment undergo dynamic changes in tissue organization, resulting in substantial changes in mechanical properties, such as increased extracellular matrix (ECM) stiffness and interstitial pressure ^15,16^. Importantly, such changes in the topological, biochemical, and mechanical properties of the microenvironment are known to alter cell states and to promote cancer aggression ^17–20^. Another highly pervasive feature of human cancers is the dysregulation of nuclear shape and size ^21,22^. Indeed, nuclear shape abnormalities have been utilized for decades as robust cancer diagnostic tools, suggesting that malignant transformation and phenotypic heterogeneity strongly associates with and may require alteration in nuclear organization. However, the precise role of nuclear architectural alteration in cancer onset and progression remains poorly understood, as does its relationship to the emergence of diversification of cellular and acellular cancer heterogeneity.

Here we develop a machine learning-based image analysis paradigm to access CRC architectural diversity and nuclear morphology and map the relationship between these features with cell state evolution and disease outcome. We find that tumor architecture, along with nuclear shapes, are highly prognostic, predictors of disease outcome. Combining the image analysis pipeline with spatial transcriptomics reveals that distinct tumor architectures correlate with nuclear shapes and specific cell states, and that architectures and states co-evolve within individual CRC tumors. Mechanistic studies with patient organoids reveal crosstalk between architecture and diversification of cell states, where tumor fragmentation and the associated mechano-osmotic stress promotes transition of classic LGR5-positive stem cells into an Annexin A1 (ANXA1)-positive fetal-like state, whereas ECM composition and crosstalk with stroma promotes epithelial-to-mesenchyme transition (EMT). Collectively these studies uncover a dynamic feed-forward loop between tissue architecture, nuclear shape and cell states that drives poor outcomes in CRC patients.

## Results

### Nuclear shape and tissue architecture are prognostic indicators in CRC

To investigate the relationship between changes in nuclear morphology of cancer cells, tissue architecture alterations, and disease outcome we obtained pathologist-curated tumor microarrays (TMAs) containing samples from a total of 1342 CRC patients with complete clinical information (Supplementary Fig. 1a). Samples consisted of TMA punches from primary tumor resection samples with paired tumor core (TC; 1825 samples) and invasive margin (IM; 1782 samples) locations for each patient (Fig. 1a, Supplementary Fig. 1b). Slides were stained for epithelial cell-cell adhesion molecule cadherin (panEPI) to identify tumor cell shapes and to separate tumor from stroma, as well as 4’,6-diamidino-2-phenylindole (DAPI) to mark all nuclei (Fig. 1a; Supplementary Fig. 1c). Nuclei were segmented at single cell resolution using a custom cellpose2.0 model (>25 million cells; ^23^) after which a panel of quantitative descriptors of nuclear shape, size, and deformation (area, circularity, solidity, aspect ratio, and EFC ratio ^24^) were extracted (Fig. 1b).

**Figure 1.**
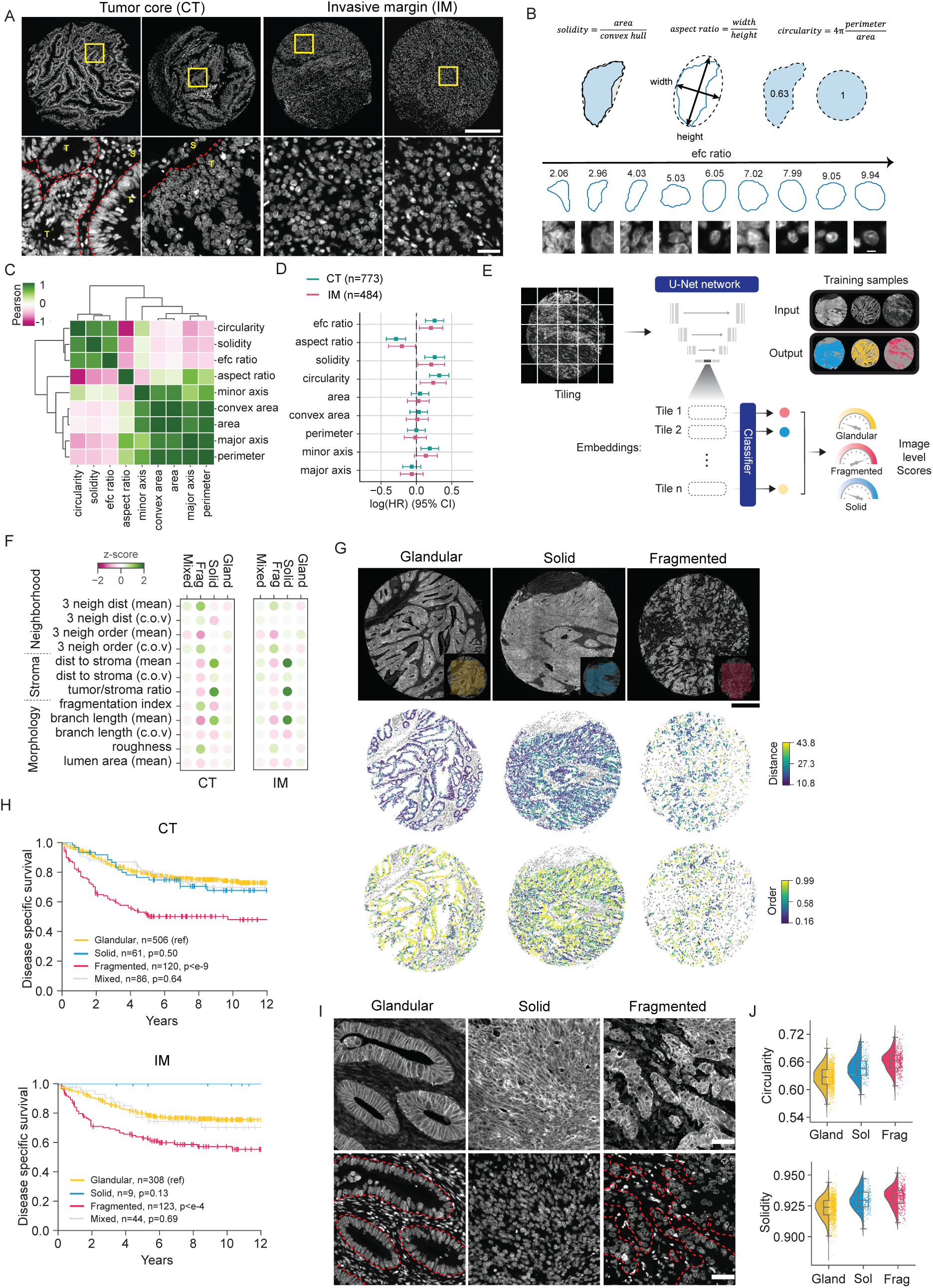
Nuclear shape and tissue architecture are prognostic indicators in CRC. **(A)** Representative images of nuclear staining (DAPI) of CRC TMA samples from different tumor core (CT) and invasive margin (IM) regions. Yellow rectangles mark areas for zoom-ins (lower panel). Separation of tumor and stroma compartments marked by red dashed line (T=tumor, S=stroma). Scale bars 500 µm (top row), 25 µm (bottom row). **(B)** Schematic description of the nuclear morphometry parameters extracted from the nuclear segmentation. **(C)** Pearson’s correlation matrix of nuclear morphometry features across all single cells from both CT and IM. **(D)** Multivariate Cox Proportional hazard model analysis of patient averaged nuclear morphometrics for disease specific survival. The x axis shows the log hazard ratio with 95% confidence interval. Note that the model includes other clinical covariates whose hazard ratios are not shown. The analysis has been conducted separately for the tumor core (blue) and the invasive margin (pink). Nuclear circularity is the strongest indicator of poor survival. **(E)** Tumor architecture classification pipeline. TMA cores with homogeneous architecture were used to train a U-Net and generate feature embeddings. Each core was then subdivided into smaller tiles, which were classified locally. The dominant (>50%) classification determined the overall architecture type. **(F)** Colored dot plots showing average values of stromal, tumor cell neighborhood, and tissue morphology descriptors across different architectures. The left plot shows measurements for the tumor core, and the right plot represents the invasive margin. **(G)** Representative TMA cores with homogeneous architectures and corresponding predictions (top row), projection of the three closest neighbors’ distance (middle row), and relative alignment (bottom row). **(H)** Kaplan-Meier curve of disease-specific survival across binned fragmentation scores from CT (top panel) and IM (bottom panel). Note poor survival of patients with high fragmentation scores in CT and IM. **(I, J)** Representative images of pan-cadherin and nuclear staining (DAPI) (I) and quantification of corresponding nuclear shapes (J) of CRC TMA samples from three architecture groups. Note increased nuclear circularity and solidity in fragmented tumors (n=1402 cores; scale bars 50 µm).

To identify the set of most informative, non-redundant parameters of nuclear morphology, we computed Pearson correlation coefficients of all parameters across single cells and clustered the parameters using hierarchical clustering (Fig. 1c). We identified two main modules of shape parameters. The first module was composed of descriptors capturing deformation or irregularities of the nucleus. EFC ratio and solidity captured deformation from perfect oval or round shape nuclei at multiple spatial scales, whereas circularity and aspect ratio mainly captured larger scale deformations (Fig. 1b, c). The second module consisted of descriptors capturing changes in nuclear size, including area and minor/major axis ratio (Fig. 1c). Each morphological descriptor was then averaged by patient and tumor region (tumor core/invasive margin) after which univariate and multivariate Cox hazard proportional models, incorporating age, stage, tumor localization and Immunoscore (a pathology-based prognostic assay quantifying T cell subsets (CD3/CD8) in tumor cores and invasive margins; ^25^), were fitted to each descriptor independently to analyze their association with survival (Fig. 1d, Supplementary Fig. 1d). This analysis revealed that nuclear deformation descriptors, rather than nuclear size descriptors, were associated with differential survival risk. Stage II and III patients with average rounder and smoother nuclei showed lower survival rate (Fig. 1d; Supplementary Fig. 1d). Notably, enhanced nuclear circularity significantly correlated with increase in tumor stage and was associated with survival in late-stage patients, whereas such association was not observed in early-stage cases (Supplementary Fig. 1e-g). Collectively, these analyses suggest that nuclear shape correlates with tumor stage and that circular nuclei associate with poor prognosis in stage II-III but not in stage I patients.

The surprising association between round nuclei, advanced tumor stage, and poor outcome led us to ask whether the morphology of single nuclei is reflective of local tissue architecture. To this end we first formalized classification of tumor architectures across patient samples. Careful histopathological classification of panEPI signal identified 5 predominant and quantifiable architecture types. Tumors with organization resembling healthy colonic glands were termed “glandular”, defined by well-polarized and tightly packed glandular structure. Glandular-like architecture with the emergence of branches and small stromal infiltrations were termed “glandular-infiltrated”. Glands with minimal stromal contact, loss of polarity, and densely packed architecture were termed “solid”. An intermediate phenotype between solid and glandular was termed “glandular dense”. Gland-like structures that exhibited detaching cell clusters were termed “moderately fragmented”. The most poorly-differentiated glands with high degree of stromal infiltration and a completely disrupted continuous gland-like architecture were termed “fragmented”.

After visually characterizing this spectrum of tissue architectures, we established a quantitative workflow to systematically classify the architectural distributions within each TMA sample into 3 main groups: glandular (consolidating glandular/glandular dense and glandular-infiltrated subtypes), solid, and fragmented (consolidating moderately and highly fragmented subtypes). To achieve this, we trained a U-Net convolutional neural network on samples with homogeneous architectures in order to generate feature embeddings (Fig. 1e). To access heterogeneity, individual TMA cores were then divided into smaller regions (tiles), and a classifier was trained using the feature embeddings to predict the architecture type for each tile (Fig. 1e, Supplementary Fig. 1h-j). The proportion of classified tiles within each category was then used to derive an overall architecture score for each patient. Specifically, the architecture classification contributing >50% of tiles was defined as the dominant architecture type for each core. Patients whose tiles did not exceed this threshold for any single architecture were labeled as “mixed.”

Having classified morphological features of tissue architectures (Fig. 1e-g) we then asked whether tissue architecture was associated with tumor stage and if it predicted patient outcome. Intriguingly, we found that architecture type both within the tumor core and at the invasive margin was highly predictive of patient outcome as measured by Kaplan-Meier analysis of disease-specific survival (DSS; Fig. 1h) and multivariate Cox proportional hazard analysis (Supplementary Fig 1k). Specifically, patients with dominant moderately or highly fragmented architectures exhibited least favorable prognosis (Fig. 1h). These cores were slightly more common in stage III and IV patients (Supplementary Fig. 1l), prompting us to ask if tumor architecture remained predictive within stage II and III patients. Kaplan-Meier analysis confirmed that tumor architecture retained its predictive value in those subgroups (Supplementary Fig. 1m). We next investigated whether patients with fragmented tumor cores also exhibited fragmented invasive margins. Nearly all patients with fragmented cores did indeed show fragmented margins; however, the reverse was not the case (Supplementary Fig. 1n). We also observed that invasive margins, in general, displayed a significantly higher degree of tissue fragmentation, and that a substantial proportion of tumors classified as solid or glandular in the core nevertheless presented with a fragmented invasive margin (Supplementary Fig. 1n-o). Collectively, these findings suggest that tumor fragmentation originates at the invasive margin and spreads to the tumor core.

To contextualize these architecture classifications within established histopathological criteria, we compared our quantification of fragmentation with the histopathological assessment of tumor budding ^26^. Tumor budding is considered a clinical marker of aggressiveness and is assessed at the invasive front of the tumor. High histopathological budding grades (BD2 and BD3) were more frequently observed in patients with fragmented architectures, indicating a partial correspondence between structural disintegration pattern and budding morphology (Supplementary Fig. 1p). Notably, even among patients with low budding scores (BD1), the presence of a fragmented architecture remained predictive, suggesting that the systematic quantification of fragmentation captures additional morphological and potentially biological features of invasion that are not fully encompassed by conventional budding grading (Supplementary Fig. 1q).

We hypothesized that cell-intrinsic nuclear morphological properties were reflective of, or resulting from evolving tissue architectures. Indeed, quantification of the relationship between nuclear shape and architecture revealed that tumor regions of patients with favorable prognosis and glandular or solid architectures, were enriched with elongated nuclei that had lower solidity (more irregular) and smaller nuclear areas. In contrast, regions with poor prognosis, enriched in both types of fragmented architectures, were dominated by nuclei that were larger, rounder, and more uniform in shape (Fig. 1i, j; Supplementary Fig. 1r).

Collectively these analyses revealed that tumor architecture within tumor cores is a strong prognostic indicator with fragmented architectures that associates with reduced patient survival. Further, nuclear shape predicts disease outcome in a manner that is dependent on tissue architecture. Round nuclei are associated with poor outcome, most likely due to their association with fragmented architecture.

### Tumor architectures correspond to transcriptional signatures of stem cell states and epithelial to mesenchymal transition

To mechanistically dissect the relationship between nuclear shape, tissue architecture and patient outcome, we proceeded to analyze the molecular signatures associated with these features. To this end we performed spatial transcriptomics (GeoMx DSP, NanoString/Bruker) analyses on a set of eight whole tumor resection samples derived from the same patient cohort used for the initial analyses (Fig 2a). For each patient, we selected 10-50 regions of interest that represented the full spectrum of architectural diversity of the given tumor (Fig. 2b) and separately profiled the transcriptomes of the tumor epithelium and its surrounding stroma (Supplementary Fig 2a). We then generated molecular clusters using Leiden clustering, defined by specific transcriptomic profiles (Fig. 2c). As expected, the transcriptomes were highly patient-specific. Interestingly, within each patient the transcriptional clusters were spatially organized into distinct domains according to distribution of tumor architectures (Fig. 2b-d; Supplementary Fig. 2b-f). This suggested that architectures could be associated with specific molecular signatures, reflecting distinct cell states. We thus assessed whether signatures of known cell states were enriched in each architecture type. Using previously published gene sets of classic, LGR5-positive crypt base columnar stem cells (CBC) and LGR5-negative regenerative/fetal-like stem cell populations ^8,14,27–29^ we discovered that moderately and highly fragmented architectures that were associated with poor disease outcomes were enriched in regenerative/fetal-like transcriptional programs, while the classic CBC signatures were enriched within glandular architectures with favorable outcomes (Fig. 2e,f; Supplementary Fig. 2 g, h).

**Figure 2.**
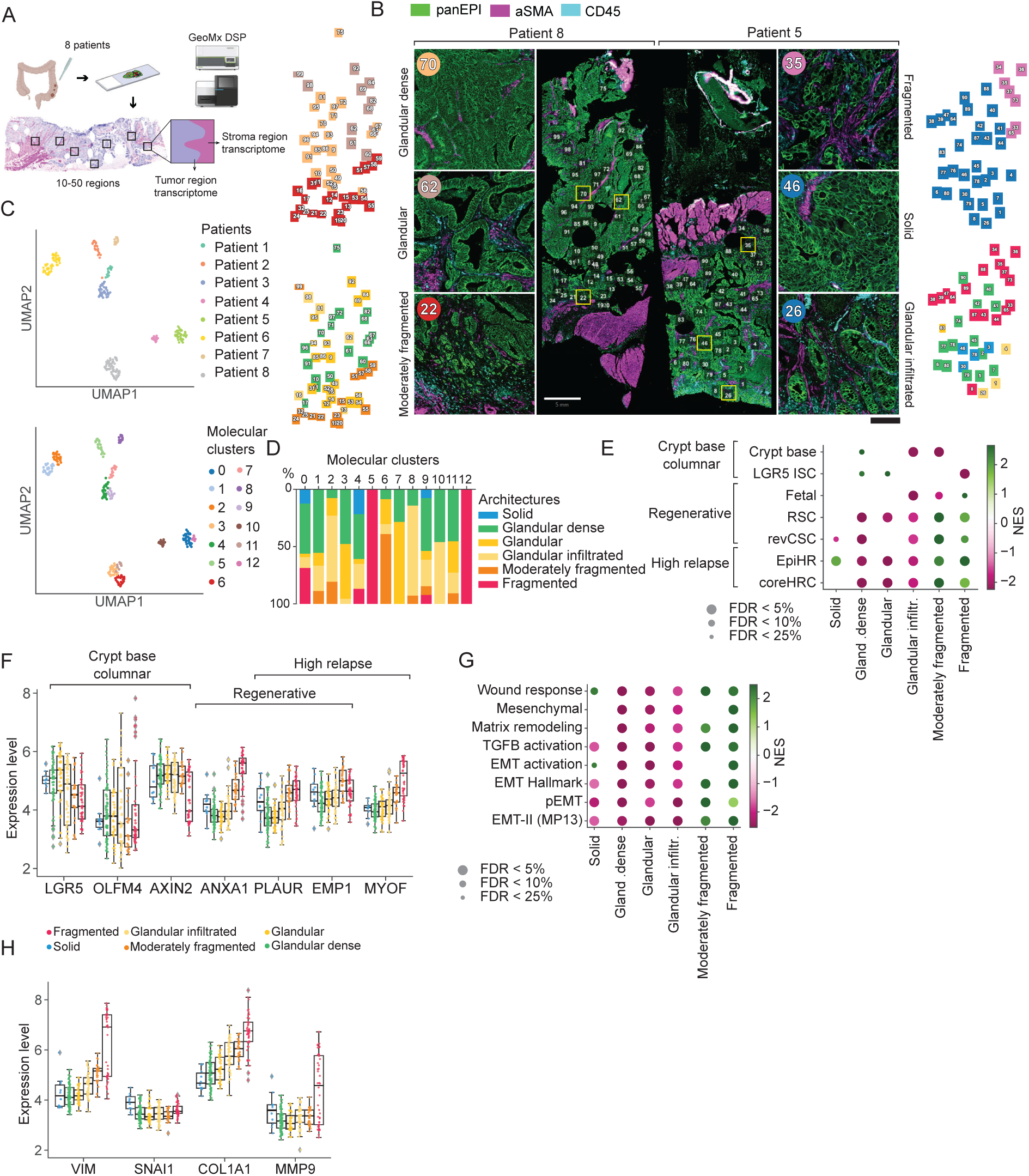
Tumor architectures correspond to transcriptional signatures of stem cell states and epithelial to mesenchymal transition. **(A)** Schematic illustration of experimental design. **(B)** Representative immunofluorescence images of pan-epithelial (panEpi), α-SMA and CD45 staining of whole patient tumors to identify tumor and stroma for selection of regions of interest (marked by numbered rectangles). Examples of two patients (patient 8 and patient 5) are shown. Numbers indicate the corresponding zoomed regions, and colors indicate the corresponding molecular clusters. **(C)** Leiden clusters of all patient samples annotated by patient identity (upper panel) or molecular identity (lower panel). **(D)** Distribution of molecular clusters across tumor architectures. **(E)** Gene set enrichment analysis of stem cell marker genes across tumor architectures. Note enrichment of CBC marker genes in glandular architectures and regenerative stem cell and high relapse gene signatures in moderately/highly fragmented architectures. **(F)** Quantification of selected key stem cell marker gene expression levels across the tumor architectures. **(G)** Gene set enrichment analysis of EMT marker genes across tumor architectures. Note enrichment of EMT marker genes in moderately/highly fragmented architectures. **(F)** Quantification of selected key EMT marker gene expression levels across the tumor architectures.

Another clinically relevant feature of malignant cells is the acquisition of mesenchymal features by the epithelial cells via the process of epithelial-to-mesenchyme transition (EMT). Similarly to classic to fetal-state intestinal stem cell reprogramming, EMT is a necessary process both during development and in adult tissue regeneration during wounding and cancer cells utilize EMT programs to escape primary tumor sites ^30^. We found that moderately/highly fragmented architectures were strongly enriched in a number of EMT signatures (Fig. 2g, h, Supplementary Fig. 2i). Collectively, these analyses reveal a clear relationship between cellular plasticity and tissue architectures in CRC with glandular and solid architectures associated with the CBC state and fragmented architectures associated with fetal/regenerative stem cell states and EMT.

### Co-evolution of cellular states and tissue architectures

The tight association of cell states and tissue architecture suggested that similarly to cell state transitions, tumor architectures might display dynamic transitions during cancer progression. Given that moderately/highly fragmented patterns were enriched in both EMT and regenerative stem cell states, we next explored if the same cells possess both regenerative stem cell and EMT features or constitute distinct populations. To determine the localization and distribution of the identified cell states within the tissue, we integrated spatial transcriptomics with multiplex immunofluorescence at single cell resolution using a panel of markers: ANXA1 (regenerative cell state), and Vimentin, Snail1, and E-cadherin to pan-cadherin ratio (EMT state). We first asked where the two stem cell states, classic and regenerative/fetal, were localized within the cancer tissue and how they were spatially distributed. In agreement with the transcriptomic data, ANXA1, a marker of regenerative/fetal stem cell states, was highly enriched in fragmented regions (Fig 3a-c, Supplementary Fig. 3a).

**Figure 3.**
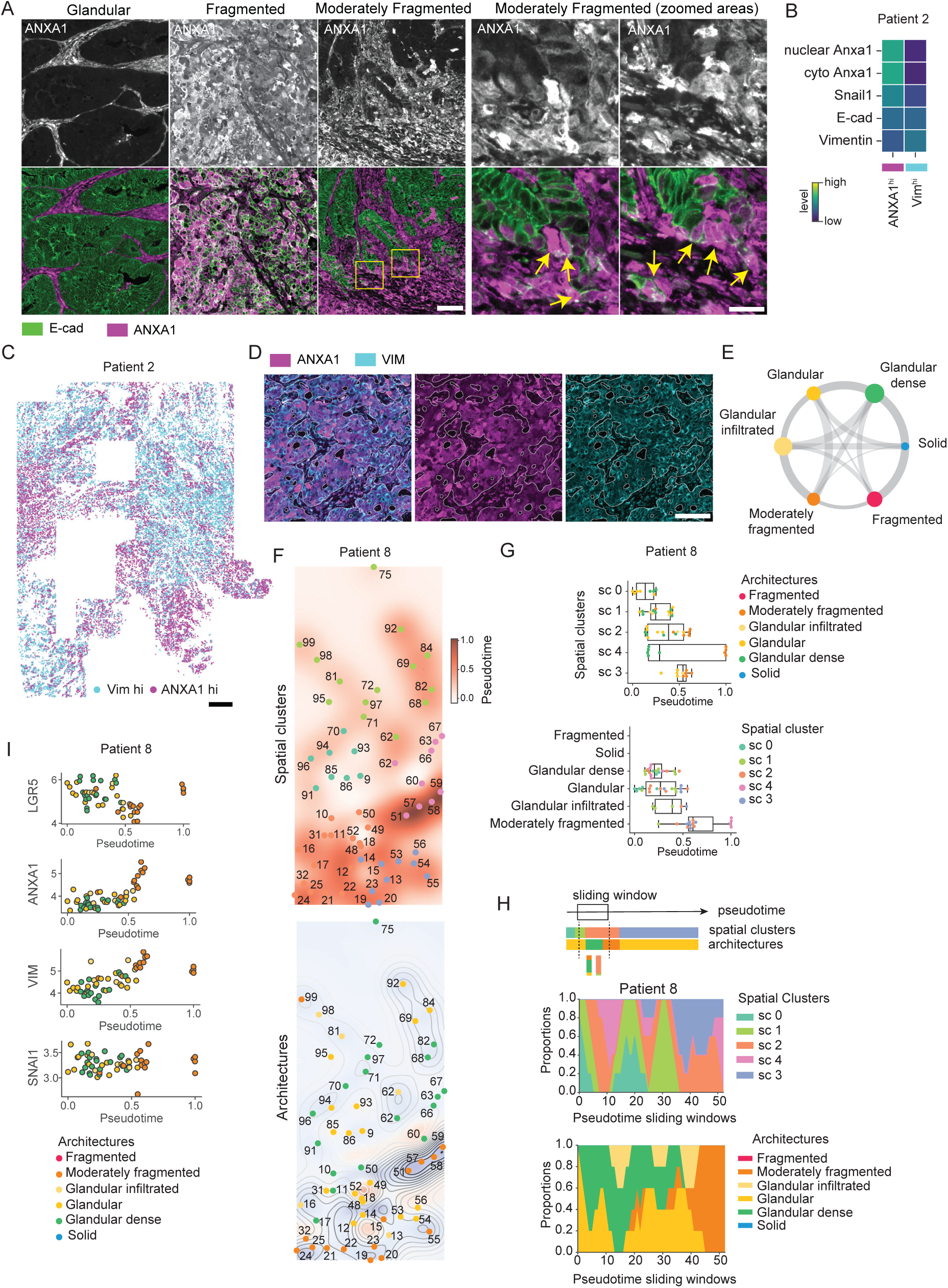
Co-evolution of cellular states and tissue architectures. **(A)** Images of E-cadherin (E-cad) and ANXA1-stained tumor biopsies from glandular fragmented and moderately fragmented regions. Note absence of ANXA1-positive cells in glandular regions, abundant ANXA1-positive cells in fragmented regions and the presence of ANXA1-positive cells at the leading edge of moderately fragmented structures (zoomed areas). Scale bars 100 µm, 30 µm (zoomed areas). **(B)** Average values of ANXA1, VIM, E-cad, and Snail1 across single-cell clusters. Note the inverse correlation between ANXA1 and EMT markers. (patient 2). **(C)** Spatial projection of ANXA1-high and VIM-high cells across the tumor (patient 2). **(D)** Representative immunofluorescence images of ANXA1 and VIM-stained tumor biopsies from fragmented regions of patient 2 from panel (A). Note lack of co-localization. Scale bars 150 µm. **(E)** PAGA trajectory inference of transitions of tumor architectures based on transcriptional profiles. Chord width reflects inferred transition probability; dot size corresponds to the number of transcriptomic regions sampled per architecture. **(F)** Representative pseudotime analyses of single patient tumors. The continuous pseudotime field (0–1) interpolated across the tissue by Gaussian Process Regression on ROI centroids is shown as heatmap. Each ROI is marked by a dot at its centroid, colored by architecture, with its ROI ID in white. The colorbar indicates the pseudotime scale (top panel); Pseudotime values are displayed as grayscale contour lines (light gray = early pseudotime; dark gray = late pseudotime). Each ROI is marked by a dot at its centroid, colored by spatial cluster, with its ROI ID in white. Regions where contours radiate outward (divergence peaks) denote sources of transcriptional progression (origin zones). Regions where contours converge (divergence) denote sinks, where the trajectory culminates (bottom panel). Note that pseudotime progression follows tissue architecture from glandular to moderately fragmented regions and aggregate within specific spatial clusters. **(G)** Boxplots showing the distribution of pseudotime values. Values are grouped by spatial clusters and colored by architecture (top); values are grouped by architecture and colored by spatial clusters (bottom). Note correlation of pseudotime with spatial clusters and architectures. **(H)** Changes in spatial clusters (top) and architecture types (bottom) along pseudotime, profiled within consecutive sliding windows. Note that distributions are relatively homogeneous at early and late pseudotime points, while intermediate segments exhibit more complex transitions, indicating a non-linear progression. **(I)** Selected marker gene expression projected across the pseudotime. Note anticorrelated evolution of LGR5, ANXA1 and VIM along the pseudotime.

As fragmented architectures showed enrichment for both regenerative/fetal-like and EMT states, we next examined their relationship. Interestingly, ANXA1 was enriched in cells located at the leading edge of the moderately fragmented regions at tumor-stroma boundary that exhibited EMT features, suggesting that regenerative stem cell state can co-exist EMT state within the tumor peripheral neighborhoods (Fig. 3a, Supplementary Fig. 3c). However, within fragmented regions these states appeared to be separated as cells exhibiting high levels of the EMT marker vimentin were distinct from those with elevated levels of ANXA1 and formed spatially segregated populations. In contrast, Snail showed similar expression levels in both populations (Fig. 3a-d; Supplementary Fig 3a, b, d). Interestingly, the expression level of Snail was also comparable between fragmented and glandular and moderately fragmented architectures (Supplementary Fig. 3a, b), which aligns with the idea that EMT represents a continuum of cell states, rather than an on/off-switch ^31^. Collectively, these analyses indicated that ANXA1-positive and -negative cell states separated into spatially segregated and distinct architecture modules. In contrast, ANXA1- and vimentin-high states were present in close spatial proximity and while exhibiting some partial overlap are mostly mutually exclusive, indicating potential plasticity between regenerative stem cell and vimentin-positive EMT states.

Since each tissue architecture type exhibited a distinct transcriptional signature with potential plasticity, we leveraged trajectory analyses of the spatial transcriptomes to test the most probable routes of architecture transitions during CRC progression. These analyses revealed that glandular architectures are statistically more likely to transition into moderately fragmented, whereas solid-like tissue architectures are highly likely to transition into fragmented architectures (Fig. 3e). Conscious of the fact that tumor architecture transitions could be influenced by an imbalance in their distribution across patients, we performed pseudotime analyses of the spatial transcriptomes from individual patients with heterogeneous architecture distributions. For each patient, we used a glandular region (as the most unperturbed architecture) in the tumor center as the reference point (time 0) and clustered the regions into spatial groups defined by nearest-neighbor to account for the possibility that similarities between regions could be confounded by their spatial proximity. We consistently observed that both the architecture and spatial cluster distributions within a given similarity (or pseudotime) window were relatively homogeneous near both the starting and end points across the three patients (Fig. 3f-h; Supplementary Fig. 3e). In addition, key markers of CBC, regenerative and EMT states exhibited steady increase or decrease along the pseudotime trajectory (Fig. 3i). Collectively, these analyses indicate that transitions between different architectures are accompanied by corresponding changes in cell states, suggestive of co-evolution of tissue structure and cellular states.

We then asked whether this potential co-evolution of cell states and architectures reflected the emergence of the distinct nuclear morphologies within the specific architectures. Indeed, consistent with our TMA analyses, single cell segmentation and quantitative analysis of nuclear morphologies revealed that glandular architectures were associated with nuclear elongation while fragmented architectures associated with an increase in nuclear size and circularity (Supplementary Fig 3f). Collectively these data showed that cell states and tissue architectures co-evolve within single patients, and that regenerative stem cell- and EMT-like states associate with fragmentation that emerges from a solid-like, CBC enriched architecture. We further observe local diversification of a potentially initially mixed regenerative stem cell /EMT state into distinct, mutually exclusive regenerative stem cell and EMT states.

### Stem cell and EMT states are controlled by distinct microenvironmental cues

To understand the mechanism that drives architecture-linked cell state diversification, we turned to disease modeling using patient-derived, whole-genome sequenced APC/TP53-mutant CRC organoids. We first asked if a change in architecture can promote the transition from a classic CBC to regenerative stem cell state. To this end we selected two independent patient-derived CRC organoid lines that, based on a shallow single cell RNAseq screen across organoid lines, had high levels of classic LGR5/OLFM4 stem cells and low levels of ANXA1/PLAUR regenerative cells (Supplementary Fig. 4a, b). We plated these two lines onto engineered colonic sheets where cells could grow unrestricted on flat surfaces or be confined in finger-like protruding channels ^32^ (Fig. 4a). Analysis of RNA levels by in situ hybridization showed heterogenous cell phenotypes with regions enriched for ANXA1-high cells that were distinct from Lgr5-high cells (Fig. 4b). Interestingly, ANXA1-high cells were enriched at bottoms of the confined finger-like channels whereas LGR5-high cells showed no preferential localization (Fig. 4b). EMT states as marked by VIM expression were homogenously distributed within the channels with partial overlap with ANXA1-high cells (Fig. 4b). Interestingly, while both patient lines responded to the architectures, the response of Line 2 was stronger than Line 1 (Fig 4b).

**Figure 4.**
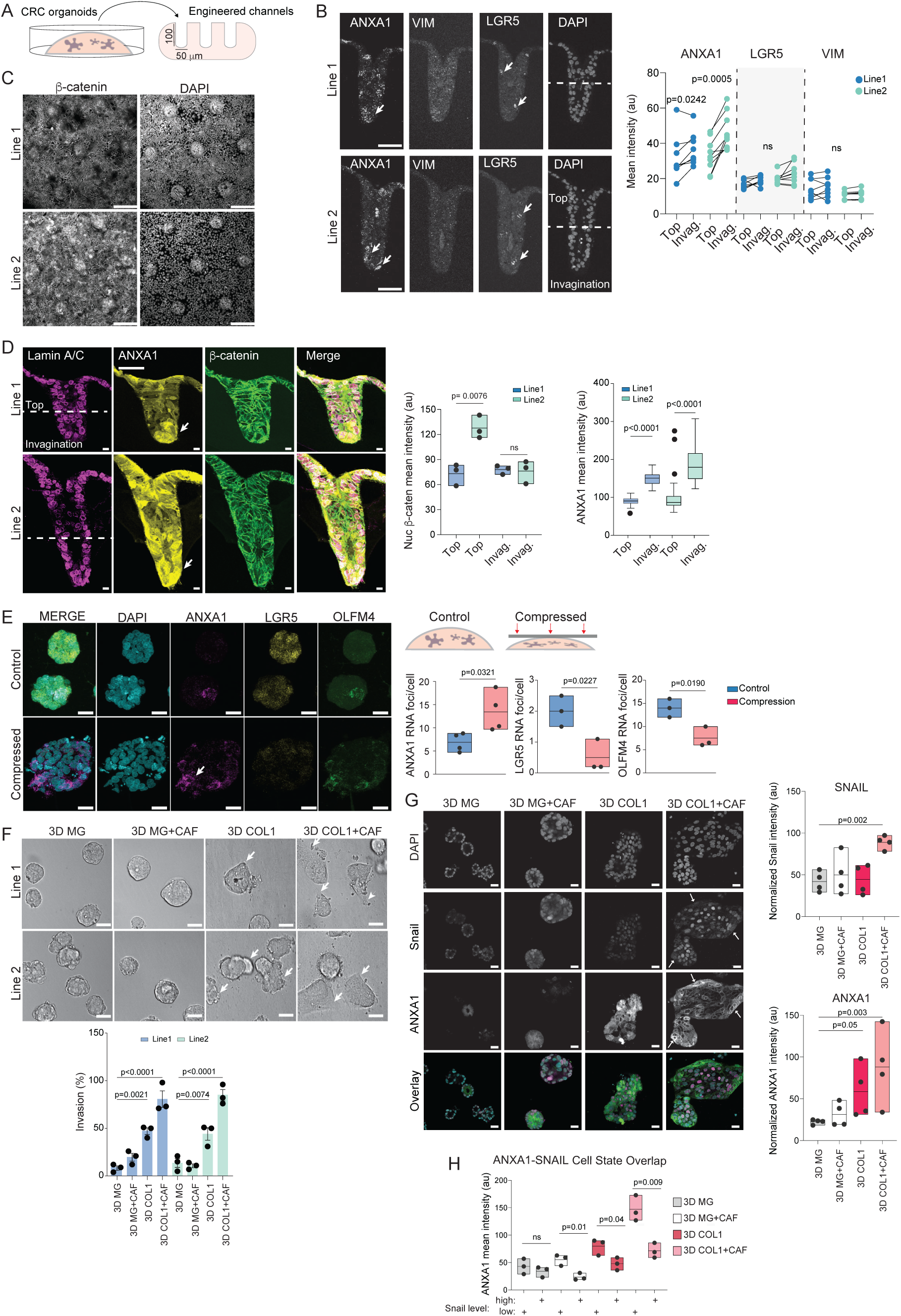
Stem cell and EMT states are controlled by distinct microenvironmental cues. **(A)** Schematic illustration of engineered colonic sheer architectures. **(B)** Representative side views and quantifications of RNAscope in situ hybridization with ANXA1, LGR5 and VIM probes of patient-derived organoids in engineered invaginating architectures. Note moderate enrichment of ANXA1-positive cells at tips of channels in Line1 and more robustly in Line 2 cells, whereas LGR5 cells show preferential localization and VIM is expressed at homogenously low levels in all cells (scale bars 50 µm; n=8 channels, Paired t-test). **(C)** Representative top views of immunofluorescence images of β-catenin-stained patient-derived organoids on flat, unconstrained surfaces. Note abundant nuclear beta-catenin in Line 2 but not in Line 1 (scale bars 100 µm). **(D)** Representative side views and quantification of β-catenin and ANXA1-stained patient-derived organoids in invaginating architectures. Lamin A/C staining is used to demarcate nuclei. Note decrease in nuclear β-catenin and enrichment of ANXA1-high cells within invaginations (scale bars 50 µm; n=3 independent experiments with >85/line/experiment, Student’s t-test). **(E)** Representative immunofluorescence images and quantification of RNA FISH for ANXA1, LGR5, and OLFM4 in controlled and compressed patient-derived organoids (**F**) Representative brightfield images and quantification of patient-derived CRC organoid lines cultured in indicated microenvironments. Note increased invasiveness in 3D collagen and 3D Collagen + CAF conditions (arrows). Scale bars 25 µm; n=3 independent experiments with >300 cells/condition/experiment, Anova/Dunnet’s (**G, H**) Representative immunofluorescence images and quantification of ANXA1- and SNAIL1-stained patient-derived CRC organoid lines cultured in indicated microenvironments. Note increased Snail expression in response to CAF conditioned medium (G) and separation of Snail and ANXA1-high cells in Collagen + CAF condition (H). Scale bars 25 µm; n=3 independent experiments with >300 cells/condition/experiment, Anova/Dunnet’s.

Examination of differences between these lines in the single cell sequencing data revealed that while both lines, when grown in Matrigel, contained high levels of classic LGR5/OLFM4-positive stem cells and low levels of ANXA1/PLAUR-positive regenerative stem cells, Line2 showed a substantially more prominent signature of WNT activity, which is a central to the regulation of stemness and EMT in the colon ^33,34^ (Supplementary Fig. 4b, c). In contrast, other molecular signatures that have been attributed to reprogramming of the fetal-like state in mice and CRC-derived human organoids, such as YAP1, SMARCC1, and SOX9 ^5^, were not differentially abundant between the two lines (Supplementary Fig. 4d). Consistent with its elevated WNT transcriptional signature, nuclear beta-catenin as readout for WNT activity was only seen in Line2 cells (Fig. 4c). Strikingly, however, within this line nuclear beta-catenin was only seen in the unrestricted, flat areas of the engineered sheets, while it was undetectable in the channels enriched in ANXA1-high cells (Fig. 4c-d). This implied that while intrinsically high WNT activity correlated with the responsiveness to tissue architecture, confinement of cells into channels suppressed WNT activity and promoted emergence of an ANXA1-positive cell state. The role of WNT activity in tuning the responsiveness to confinement was confirmed by culturing the responsive cell line in the presence or absence of WNT3a/RSPO. As expected, adding WNT into the medium enhanced the impact of confinement on ANXA1 expression (Supplementary Fig 4e)

To test if confinement indeed promoted stem cell fate transitions, we cultured the patient organoids in Matrigel, were ANXA1 levels were initially low (Fig. 4e; Supplementary Fig. 4b), and subjected them to mechanical confinement using a compression bioreactor ^35^. Indeed, confining the organoids was sufficient to trigger emergence of ANXA1/PLAUR-positive cells with corresponding suppression of LGR5/OLFM4-positive cells (Fig. 4e).

To confirm the role of architecture in a setting more closely mimicking fibrotic ECM remodeling during CRC progression^36,37^, and to understand if state changes would be associated with changes in cell behavior, we further compared organoids grown in soft Matrigel to a confining environment generated by stiff 3D Collagen. Indeed, this switch was sufficient to trigger moderate invasive behavior (Fig. 4f). To test if soluble microenvironmental signals could collaborate with architecture-induced confinement, we supplemented the stiff-collagen cultures with conditioned medium from patient-derived CAFs. While the conditioned medium was not sufficient to trigger invasiveness in Matrigel, it strongly enhanced invasive behavior in 3D Collagen (Fig. 4f). We hypothesized that this switch in behavior could be triggered by promotion of EMT, that was not enhanced by the confining architectures alone (Fig. 4b). Indeed, CAF conditioned medium strongly stimulated expression of the EMT master regulator transcription factor Snail, specifically in the 3D Collagen conditions (Fig. 4g). Intriguingly, while the Snail and ANXA1-positive cells overlapped in the Matrigel conditions, they became spatially segregated in stiff 3D Collagen + conditioned media conditions (Fig. 4h), resembling the separation of these states at the invasive margin of patient cells (see Fig 3). A similar effect was found in compressed organoids, where the ANXA1-high cell state, triggered by compression, appeared mutually exclusive from the Snail-high cells that were not influence by compression (Supplementary Fig. 4f, g). Collectively these experiments showed that tissue architecture and stromal signals collaborate to promote fetal-like stem cell states and EMT, where confinement suppresses WNT to promote stem cell fate transitions and soluble signals enhance EMT features.

The combination of the two microenvironmental signals drive diversification of the two states and invasive behavior.

## Discussion

Identifying the mechanisms driving intratumoral heterogeneity and cellular diversity is central towards achieving improved diagnostic precision and curative treatment for CRC. Our data identifies tumor architecture and the associated nuclear shapes as strong prognostic indicators in CRC. We further demonstrate that tumor architectures and cell states dynamically co-evolve within individual patients, with fragmented architectures associated with a transition of the classic LGR5-marked stem cell state with a fetal-like regenerative stem cell state that has been associated with aggressive behaviors and poor prognosis ^8^.

We find that the power of the tumor architecture quantification is based on the co-evolutionary relationship of aggressive cell states and fragmented tumor architecture. It is well established that cellular stress triggered by chemotherapy, wounding, inflammation or mechanical damage induces phenotypically flexible cell states which, in the context of CRC, involve the reactivation of the fetal stem cell program ^29,38–42^. Additionally, the activation of TGFβ1 has been shown to drive Wnt-independence and lineage conversion of CRC-derived organoids into fetal-like state ^43^. On the other hand, fibrosis-driven increased stiffness of the tumor stroma has been shown to drive aggressive behaviors in a large number of solid tumors ^44^. Intriguingly, our study implicates compressive forces imparted by the stiffened extracellular matrix which distorts tissue geometry and induces tumor fragmentation to promote the transition from the classic to the fetal-like stem cell state. This fetal-like stem cell state first emerges as a hybrid state with EMT features. Additional signals from the tumor stroma then further diversify these states into spatially closely associated but distinct EMT and stem cell states. The state diversification is associated with triggering invasive behavior, explaining the strong correlation between fragmented architectures and poor prognosis. These observations support a model where disruption of tissue architecture and inflammatory signals initiate a co-evolutionary feed forward loop that drives disease aggression.

The image analysis paradigm developed in this study provides an inexpensive and easily implemented patient biopsy stratification strategy, based on immunostaining of nuclei and an epithelial membrane marker coupled to machine learning-based image analysis, to classify tumor architecture and cell shapes that may aid in predicting patient outcome. Importantly, our work suggests that quantitative analysis of tumor heterogeneity using architecture as proxy may be the key to diagnostic precision.

## Methods

### Patient samples

The study uses a previously described cohort ^45^ comprising 1343 patients who underwent resection for colorectal cancer at Central Finland Central Hospital between January 1st, 2000 and December 31st, 2015 and had adequate tumour samples available. Formalin-fixed and paraffin-embedded (FFPE) tumor samples were obtained from the pathology archives of the Central Finland Central Hospital Nova, which covers all colorectal cancers diagnosed in Central Finland (the population of the area-averaged around 270,000 during the study period). Associated clinical data were collected from clinical patient records by study physicians. In addition, all tumors were previously re-evaluated for disease stage, tumour grade (WHO 2019 criteria), MMR status, *BRAF* V600E mutation status, and the presence of lymphovascular invasion ^46^. Analyses of this cohort were approved by the Regional medical research ethics committee of the Wellbeing services county of Central Finland (Dnro 13U/2011, 1/2016, 8/2020, and 2/2023), the Finnish Medicines Agency (Fimea, FIMEA/2023/001573). The need to obtain informed consent from the study patients was waived (Dnro FIMEA/2023/001573). Final TMA blocks of duplicate 1.2 mm cores were made in TMA Grand Master (3D Histech). For spatial transcriptomics selected full tumor samples from the same cohort were used. The authors affirm that the study was conducted following the rules of the Declaration of Helsinki of 1975, revised in 2013.

### Immunofluorescence staining and imaging of patient samples

Immunofluorescence staining and imaging of TMAs was performed as previously described ^47^ for pan-cadherin antibodies to label the tumor cells and their plasma membranes and the nuclear marker DAPI. Imaging was performed using a Zeiss Axio Scan.Z1 slide scanner, with each round of staining recorded as an independent. CZI image file containing up to five fluorescent channels. For the full tumor samples, the staining was carried out on the same slides as spatial transcriptomics, following the spatial transcriptomics analyses. In order to select ROIs for sample collection for transcriptome analysis, the slides were stained with Syto 13, PanCK (AF532), CD45 (AF594) and aSMA (F647) to visualize nuclei, epithelial cells, immune cells and stroma, respectively. Imaging of the staining was done on the DSP instrument using 50, 200, 200, 200 ms on the FITC, CY3, Texas Red and CY5 channels, respectively. After quenching the signal, the slides were restained for indicated markers and the signal was imaged on the DSP instrument. The single channel images were loaded to HALO vs 3.6 image analysis software for visualization, registered based on the nuclear signal and fused to generate a composite image with 8 markers (including the nuclei).

### Image analysis pipeline for patient samples

#### Preprocessing and segmentation

Preprocessing and segmentation were performed essentially as described earlier ^48^. Briefly, TMA cores were de-arrayed using built-in tools from QuPath (QuPath 0.5.1; ^49^) Cores exhibiting folding artifacts, blurriness, or limited tumor content were excluded from analysis. After preprocessing, Nuclei were segmented from DAPI signal using custom-trained Cellpose 2.0 models ^50^, separately optimized for the TMA and whole-slide image (WSI) datasets. Tumor and stromal compartments were identified using a multi-label U-Net trained on 30 representative cores. Pixel-level annotations were generated manually in ilastik (version 1.4.0; ^51^) based on pan-epithelial (pan-EPI) staining.

#### Architecture classification of TMAs

A second multi-label U-Net was trained on 196 cores selected for their homogeneous architectures. Training masks were generated from the tumor-stroma segmentations obtained in the previous step, with tumor regions re-annotated according to their specific architectural subtype. Each core was then subdivided into 312×312 px tiles, and features were extracted from the bottleneck layer of the second U-Net for each tile. To refine classification, a multilayer perceptron (MLP) with two hidden layers (200 and 100 neurons), tanh activation, and the Adam optimizer was trained on 950 tiles (202 fragmented, 321 glandular, 120 solid, 149 stroma, and 157 edge/background). Performance was evaluated using stratified five-fold cross-validation. Finally, tile-level predictions were aggregated per patient: if one architecture exceeded 50% of the tiles, it was labeled as the “dominant” architecture; otherwise, the sample was classified as “mixed.”

#### Morphological features of tissue architectures

Branch lengths were quantified from the binary mask of the tumor-stroma mask by first skeletonizing the mask using the skimage.morphology.skeletonize. Small spurious branches shorter than 5 px were removed via the prune_skeleton function. Global contour roughness of the tumor boundary was assessed by extracting all contours from the binary mask (find_contours at level 0.5) using skimage.measure.find_contours. Contours with fewer than 5 points were discarded. The rest were smoothed (Gaussian σ = 1) to reduce pixel noise. Pointwise curvature κ for a smoothed contour (x(s),y(s)) was computed by finite differences:

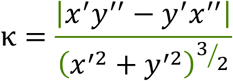

where primes denote derivatives with respect to the contour parameters. Absolute curvature values were summed over all contours and normalized by the total number of contour points

*N* to yield mean roughness per unit length.

Lumens within the binary mask were detected by filling (binary_fill_holes) and subtracting the original mask. The resulting lumen mask was labeled, and each lumen (in pixels) was measured via skimage.measure.regionprops.

3-nearest-neighbor distances were computed by constructing a KD-tree of all cell centroids and querying, for each centroid, the Euclidean distances to its three closest neighbors.

Stroma proximity was determined by applying a Euclidean distance transform to the binary stroma mask and sampling the resulting distance map at each centroid to obtain its minimal distance to stromal tissue.

Tumor–stroma ratio was calculated by measuring the total pixel areas of the tumor and stroma masks and computing the ratio of tumor mask area to stroma mask area. Fragmentation index was computed by first constructing a k-nearest-neighbor (KNN) graph of the cell centroids to derive spatial communities (clusters). The fragmentation index was then calculated as the number of clusters divided by the total pixel area of the tumor mask.

#### Whole slide image (WSI) analysis

Whole-slide image (WSI) immunostainings were registered using the warpy workflow in ImageJ and QuPath. WSIs were subsequently divided into non-overlapping tiles of 1000 × 1000 pixels and selected for downstream analysis in a manner analogous to tumor microarrays, including tumor/stroma classification and nuclear segmentation. For each nucleus, the average nuclear intensity of markers (ANXA1, Snail1) was quantified. For cytoplasmic markers (E-Cadherin, ANXA1, VIM), the nuclear contour was dilated by 5 pixels to approximate the cytoplasmic region, from which the average cytoplasmic intensity was computed. Single-cell profiles were then clustered using the Leiden algorithm at a resolution of 0.1.

#### Spatial Coherence

For each whole-slide-image (WSI) tile we computed a spatial-coherence score that quantifies whether cells with the same label group together or segregate. First, for every cell we counted its 12 immediate closest neighbours that shared the same label and averaged this count across all cells in the tile. Next, we generated a null range by randomly shuffling the position of all cells 100 times, repeating the same neighbour counting for each shuffle; the average value from these shuffled sets was taken as the expected mean. The score was then calculated as (observed – expected_mean)/(expected_max – expected_mean). A score above zero means cells with the same label cluster together more than would be expected by chance (attraction), a value of zero corresponds to a random arrangement, and a negative value denotes spatial repulsion, where same-label cells are less likely to be adjacent than expected by chance.

### Spatial transcriptomics experiment and analyses

Spatial transcriptomics was performed using the GeoMX-DSP platform (Nanostring/Bruker). Eight stage II-III patients were selected from the TMA cohort based on presence of architecturally heterogeneous tumors. Samples were prepared following recommendations from the vendor for FFPE slides, including baking for 60 min at 60oC, dewaxing and antigen retrieval with ER2 for 20 min at 100oC, fixation in NBF (10%), incubation in NBF top, and proteinase K treatment (15 min at 37oC). These steps were performed on the Leica BondRX autostainer. The slides were incubated with the probes overnight and, after stringent washes, were stained with Syto13, pan-cytokeratin (AF532), aSMA (custom conjugated on AF647) and CD45 (AF594) antibodies to detect tumor and stromal microenvironments. Regions of interest (ROIs) (67, 61, 96 and 59 for slides 1-2, 3-5, 4-6 and 7-8, respectively) were segmented for panCK positivity to collect 82, 94, 141 and 63 areas of interest (AOIs) representing tumor and tumor-stroma boundary. After scanning, tissue sections were additionally stained with antibodies against Snail1, Ecad, ANXA1 and Vimentin and imaged using Phenoimager (Akoya Bioscences).

Quality control of the sequencing data was performed in R using GeoMx packages, following standard GeoMx QC guidelines. Segment filtering employed the following thresholds: minimum reads per segment=1000, minimum percentage of reads trimmed=80, minimum percentage of reads stitched=80, minimum percentage of reads aligned=75, minimum sequencing saturation=50, minimum negative control counts=1, maximum counts in the NTC (no-template control) well=9000.

Segments with low signal were removed based on the limit of quantification (LOQ). Gene filtering was applied by both compartment (tumor/stroma) and architecture type to preserve important genes in underrepresented regions. A 10% segment cutoff was used. Data were normalized using the third quartile (Q3) method.

#### Differential gene expression analysis

For differential gene expression, raw counts were analyzed using the PyDESeq2 Python package, with patient sample included as a covariate to control for inter-patient variability. For each gene, a Wald test was used to assess differential expression; multiple-testing correction was performed using the Benjamini–Hochberg procedure to control the false discovery rate (FDR). Genes with an adjusted p-value (q-value) below 0.05 were considered significantly differentially expressed.

#### Gene set enrichment analysis

Gene set enrichment analysis was performed using the prerank function from the GSEApy Python package with 1,000 permutations. Prior to analysis, all genes were ranked according to their effect sizes (log fold changes) derived from the differential expression analysis. The following gene sets from previous studies were included:

#### EMT gene sets

Wound response GO GP ^52^, Mesenchymal ^53^, Matrix remodeling ^54^, TGFB activation (TGFB KEGG, TGFB CORE GENES / TGFB 1 / TGFB 2) ^54,55^, EMT activation (EMT CORE GENES ^53^, EMT Hallmark ^56^, pEMT ^57^, EMT-II (MP13) ^58^.

#### Stem cells gene sets

Crypt base ^27^, LGR5 intestinal stem cells ^28^, Fetal ^8^, Regenerative ^29^, revival colonic stem celles ^39^

#### High relapse gene sets

Epithelial-specific high-risk gene set, core high-relapse cells ^59^.

#### Trajectory analysis

Trajectory analysis was performed using partition-based graph abstraction (PAGA) ^60^ with the sc.tl.paga function built into Scanpy ^61^, version 1.9.6). To prepare the data, we first selected 3,000 highly variable genes using the “seurat” flavor of sc.pp.highly_variable_genes. Next, we calculated and selected100 principal components (PCs) using sc.tl.pca. PCs were batch-corrected using Harmony (^62^, version 0.0.9) to correct for patient-specific gene expression profiles. Finally, nearest neighbors were calculated using sc.pp.neighbors (n_neighbors=30) on the corrected PCs and PAGA was run on tumor architecture types.

#### Pseudotime analysis

We calculated diffusion pseudotime (DPT^63^ using the sc.tl.diffmap and sc.tl.dpt functions from Scanpy ^61^, version 1.9.6). In preparation, for a patient sample we selected 3,000 highly variable genes using the “seurat” flavor for sc.pp.highly_variable_genes. Next, we calculated 100 principal components (PCs) using sc.tl.pca. Finally, nearest neighbors were calculated using sc.pp.neighbors (n_neighbors=30) on the corrected PCs. DPT was run using a cell in the glandular region of the tumor as “root cell”, representing its most “unperturbed” region.

### Patient-derived 3D organoid culture

The utilization of human CRC organoid cultures was approved as part of the iCAN Precision Medicine Flagship study reviewed by the HUS Ethical Committee and is executed based on a HUS research permit (4.5.2023 §38 (HUS/223/2023), a Findata data permit (THL/1338/14.02.00/2022), and MTAs with Helsinki Biobank (HBP20210170) and Finnish Hematology and Registry Biobank FHRB (12.5.2022). The underlying PANORG study collecting the samples to the iCAN Flagship was approved by the ethical committee of the Helsinki and Uusimaa hospital district and study site institutional review boards (PANORG ethical approval HUS/390/2021, HUS approval HUS/69/2022). The study participants provided a written, informed consent for the protocols. Organoids were maintained in a mixture of 60% LWRN media (originally established in ^64^) 40% human basal media supplemented with rhEGF (R&D, 236-EG, 100ng/ml); LWRN was replaced with Advanced DMEM (ThermoFisher) for “No Wnt” conditions. Organoids were subjected to single cell sequencing and selected to harbor APC and TP53 mutations for the studies here.

### Single cell RNA sequencing

Patient-derived CRC organoids were cultured in Matrigel, after which single cell suspensions were generated by incubation with Cell Recovery Solution (30 minutes at +4C; Corning) followed by digestion with TrypLE™ Express Enzyme (5minutes at 37°C; Invitrogen). After washing with ice-cold DMEM, single cells were counted using Luna-II automated cell counter (Logos Biosystems) and loaded on a microwell cartridge of the BD Rhapsody Express system (BD) following the manufacturer’s instructions. Single cell whole transcriptome analysis libraries were prepared according to the manufacturer’s instructions using BD Rhapsody WTA Reagent kit (BD, 633802) and sequenced on the Illumina NextSeq 500 using High Output Kit v2.5 (150 cycles, Illumina) for 2 × 75 bp paired-end reads with 8 bp single index aiming sequencing depth of >20,000 reads per cell for each sample.

#### Analysis

UMI, complex barcode, and sample tags were extracted from raw reads and demultiplexed using a custom script (rhapsody-demultiplex, https://github.com/Bioinformatics-Service-MPI-Munster/scrna-tools). Reads were mapped using STAR version 2.7.10a ^65^ (--soloType CB_UMI_Complex --soloCellFilter None --outSAMtype BAM SortedByCoordinate -- soloFeatures GeneFull_Ex50pAS --soloAdapterSequence NNNNNNNNNGTGANNNNNNNNNGACA --soloCBmatchWLtype 1MM --soloCBposition 2_0_2_8 2_13_2_21 3_1_3_9 --soloUMIposition 3_10_3_17 --soloCBwhitelist BD_CLS1_v2_draft.txt BD_CLS2_v2_draft.txt BD_CLS3_v2_draft.txt --runRNGseed 1 -- soloMultiMappers EM --readFilesCommand zcat --outSAMattributes NH HI AS nM NM MD jM jI MC ch CB UB GX GN). Raw counts were imported as AnnData ^66^ (v. 0.10.7) objects in Python (3.9.11). We removed low complexity barcodes with the knee plot method (1500-3000 counts per cell), and further filtered out cells with a high mitochondrial mRNA percentage (>50%). Doublets were predicted with scrublet ^67^ (v. 0.2.3). Finally, each sample’s gene expression matrix was normalized using scran ^68^ (v. 1.26.2), with Leiden clustering ^69^ input at resolution 0.5.

At this stage samples were merged with scanpy ^61^ (v. 1.10.1). For 2D embedding, expression values were cell-cycle regressed using sc.tl.score_genes_cell_cycle with G2M and S-phase gene lists from ^70^and sc.pp.regress_out. The expression matrix was subset to the 3,000 most highly variable genes (sc.pp.highly_variable_genes, flavor “seurat”). We calculated and selected the top 100 principal components (PCs) (sc.tl.pca), which served as basis for k-nearest neighbor calculation (sc.pp.neighbors, n_neighbors=30). Nearest neighbors were then used as input for UMAP layout (sc.tl.umap, min_dist=0.3). Leiden clusters were calculated using sc.tl.leiden at resolution 0.7.

### Engineered hydrogel scaffolds

#### Microstamping hydrogel topography

Hydrogel scaffolds were engineered as described previously ^32,71^. A PDMS stamp (see below) with crypt-mimmicking array of pillar topography was placed in the central top reservoir of the chip. To generate the ECM for the chip bovine TeloCol-6 Type I Collagen solution (5225-50mL, Advanced Biomatrix) was neutralized following manufacturer instructions and then mixed with Matrigel (356231, Corning) in 75/25 (v/v) ratio. After gel injection through gel-loading port, hydrogel was polymerized at 37°C, 5% CO2 (hereafter referred to as “incubator”) for 15 min. After ECM polymerization, all chip reservoirs were topped up with PBS or cell culture media. Prepared chips with hydrogels can be stored for multiple weeks in the incubator or in the +4°C fridge before use.

#### Microtopography stamps fabrication

Stamps featuring crypt-mimicking microtopography were fabricated from PDMS via replica molding of 2PP-printed models. First, arrays of colon crypt-mimicking micropillars were designed in Autodesk Fusion. Several topographies were tested and optimized, the final microtopography design consists of the slightly conical pillars, ranging from 70 to 120µm in diameter (top to bottom) and 700µm in height. The 3D model was printed using a NanoOne (UpNano) 2PP printer on glass coverslips with UpPhoto resin and a 10X 0.4 NA objective. To generate a master mold with inverted topography, the printed structures were plasma-activated and silanized overnight using vapor-phase trichloro(1H,1H,2H,2H-perfluorooctyl) silane (Sigma-Aldrich). PDMS (Sylgard 184, Dow Corning; 10:1 base-to-curing agent ratio) was poured over the master, degassed under vacuum, and cured at 80°C for 12h. The resulting PDMS replica was cut, plasma-activated, and silanized overnight as above. This PDMS negative was then used repeatedly as a mold for casting final PDMS stamps using the same procedure (Sylgard 184, 10:1 ratio, cured at 80°C for 12h).

### RNA in situ hybridization

In situ hybridization was performed using RNAscope technology (Advanced Cell Diagnostics) according to the manufacturer’s protocols. Briefly, organoids were fixed in 4% PFA for 30 min at 37°C, rinsed with PBS, dehydrated in a graded ethanol series and air dried. Slides were then rehydrated and 2 drops of hydrogen peroxide was added to each slide for 10 min. Primary antibodies were added and incubated overnight at 4°C. Slides were wshed and fix with 4% PFA for 30 min at room temperature. After PBS washes, 3 drops of 1:15 diluted Protease Plus was added to slides and incubated for 10 min, followed by PBS washes.

RNA probes were prewarmed for 10 min at 40°C, cooled down and added on slides. Slides were incubated with probes for 2h at 40°C, washed twice with wash buffer, followed by stepwise hybridization of amplifier solutions 15 min each at 40°C, followed by blocker solutions. Finally, samples were incubated with Alexa-conjugated secondary antibodies diluted in Co-detection Antibody Diluent for 30 min at room temperature. After washes and brief incubation with DAPI, slides were mounted in Elvanol and imaged.

### Mechanical compression of organoids

To mechanically compress the organoids, organoids grown in Matrigel for 5 days were replated on top of Matrigel coated 12-20mm round glass coverslips and allowed to attach for 24 hours. Post attachment, organoids were sandwiched with another 12-20mm coverslip with a weight applying 200 Pa of vertical stress (generating 45% reduction in organoid height) for 20 hours

### Statistics and Reproducibility

Statistical analyses were performed using GraphPad Prism software (GraphPad, version 9) or in R (version 2023.09.1+494). Statistical significance was determined by the specific tests indicated in the corresponding figure legends. Only 2-tailed tests were used. In all cases where a test for normally distributed data was used, normal distribution was confirmed with the Kolmogorov– Smirnov test (α = 0.05). All experiments presented in the manuscript were repeated at least in 3 independent replicates.

## Acknowledgements

We thank Claudia Ortmeier, Manuela Haustein, Julia Kolikova, Meiju Kukkonen and Hanne Ahola for technical assistance, Jason R. Spence and Michael K. Dame (University of Michigan) for advice and support on organoid cultures, the Molecular Histopathology Laboratory (MHL) at NCI, NCI/NIH for whole slide scanning, and Max Planck Institute Sequencing Core Facility for support with sequencing experiments. We thank the cancer patients, as well as the study nurses Jaana Koskinen-Alhainen, Katri Suomala and Tuija Vanhatalo for providing the valuable patient samples for the study and Dr. Juha Väyrynen (Oulu University) and Dr. Erkki-Ville Wirta (Pirha) for the curated TMA dataset. We acknowledge participants and investigators of the iCAN study. The iCAN Flagship Project is funded by the Research Council of Finland (grant number 346555), University of Helsinki, Helsinki University Hospital and industry partner Boehringer Ingelheim. This work was supported by funding from Jane and Aatos Erkko Foundation, Sigrid Juselius Foundation, Mary and Georg Ehrnrooth Foundation, Cancer Foundation Finland, Relander Foundation, and HUS and Pirha state research funding (all to TTS), the Max Planck Society, the Helsinki Institute of Life Science, Academy of Finland Molecular Regulatory Networks of Life (#330033, NucleoMech), and the Academy of Finland Center of Excellence BarrierForce (all to SAW), and Intramural Research Program of the NIH, National Institute of Diabetes and Digestive and Kidney Diseases (NIDDK) and Orion Foundation (all to YAM).

## Author contributions

FB performed all quantitative image and sequencing analyses with input from KK. SH performed *in vitro* organoid work with support from YAM, ND, and MB. ME and YAM performed multiplex immunohistochemistry work. NK and MOH performed whole slide imaging and GeoMx experiments, TP and KP provided advice on multiplexing. CH, JH, JB, J-PM, and TTS provided advice on all the clinical aspects of the study. TTS provided all tumor samples and organoids, EPE supported with organoid analyses, MA prepared tumor sections. MN, NG and ML enabled colonic sheet experiments. SAW and YAM conceived the study, designed experiments, provided resources and conceptual advice. YAM conceived and directed the study, provided resources, designed and performed experiments, and analyzed data. YAM and FB wrote the manuscript with input from all authors.

---

**Supplementary Fig. 1.**
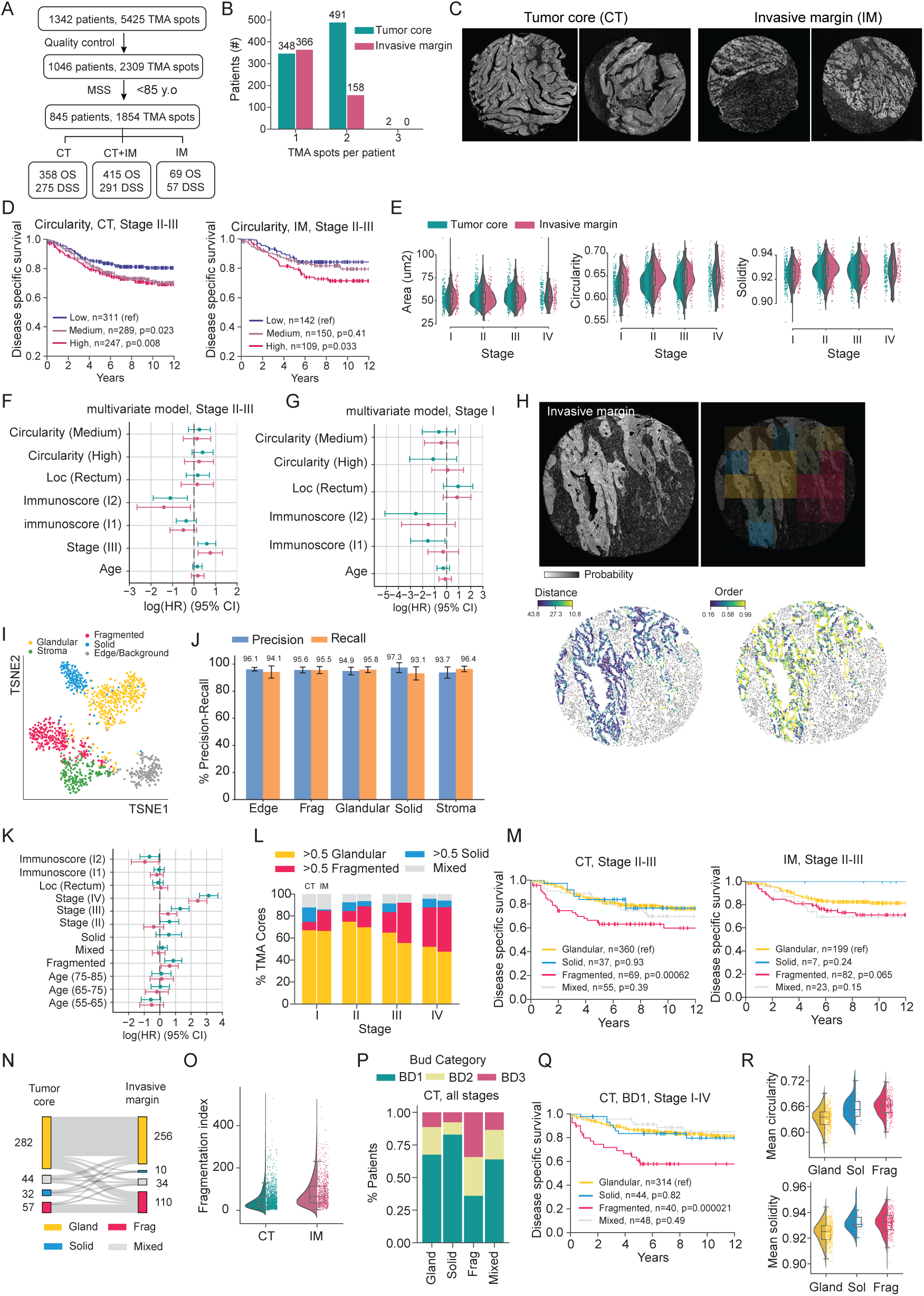
**(A)** Patient dataset and analyzed samples. **(B)** Number of analyzed TMA cores per patient for each compartment (tumor core and invasive margin). **(C)** Representative images of tumor cores and invasive margins. **(D)** Kaplan-Meier survival curves for patients stratified by nuclear circularity (high, medium, low) of tumor core (left) or invasive margin (right). **(E)** Nuclear shape (area, circularity, and solidity) per patient, grouped by tumor stage. **(F, G)** Multivariate Cox proportional hazardsmodels including the circularity shape descriptor, comparing stage II–III (F) with stage I (G) patients. Circularity is associated with poor survival in stage II–III but not in stage I patients. **(H)** Representative invasive margin sample with heterogeneous architectures: heterogeneous architecture (top left), classifier predictions (top right), projection of three closest neighbors’ distances (bottom left), and relative alignment (bottom right). **(I)** t-SNE embedding of features derived from the U-Net. **(J)** Precision and recall for architecture classification across classes, with error bars representing the standard deviation from five-fold cross-validation. **(K)** Multivariate Cox proportional hazards models incorporating Solid, Fragmented, and Mixed architecture types, with Glandular as the reference. Note that the Fragmented architecture remains significantly associated with poor survival after adjustment for other clinical covariates. **(L)** Stacked bar plot displaying the distribution of architecture types across tumor stages, with left bars representing the tumor core and right bars representing the invasive margin. **(M)** Kaplan-Meier survival curves for stage II–III patients, stratified by architecture type, with the right panel showing tumor core and the left panel showing invasive margin data. Patients with fragmented architectures exhibit poorer survival. **(N)** Sankey diagram illustrating the correspondence between tumor core and invasive margin architecture classifications, indicating that all fragmented tumor core samples also display fragmented invasive margins, although the reverse is not observed. **(O)** Distribution of the fragmentation index per TMA core, comparing values between the tumor core and invasive margin. **(P)** Stacked bar plot displaying the distribution of budding grades defined by the International Tumor Budding Consensus Conference (ITBCC, 2016) across architecture types. **(Q)** Kaplan-Meier survival curves for stage I–IV patients with low budding BD1grade, stratified by tumor architecture. Even among patients with low budding, those with fragmented architectures show significantly poorer survival, suggesting that fragmentation captures additional prognostic information beyond budding alone. **(R)** Patient-average nuclear shape descriptors (circularity and solidity) for TMA cores across different architecture types in the invasive margin.

**Supplementary Fig. 2.**
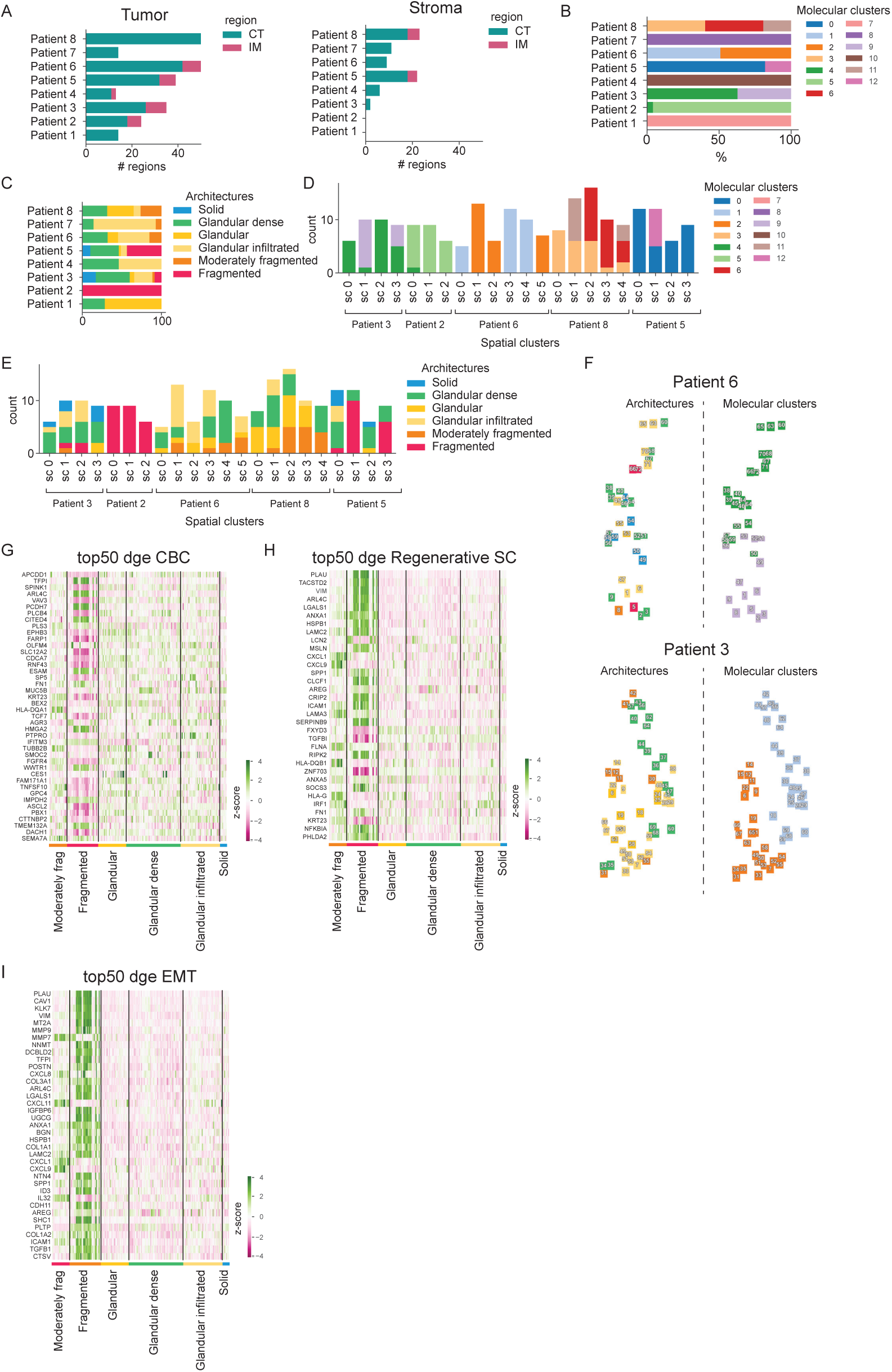
**(A)** Number of transcriptomic regions per patient, stratified by compartment (tumor core vs. invasive margin) and tissue type (tumor on left, stroma on right). **(B)** Distribution of molecular clusters across patients, demonstrating that each patient harbors unique clusters. **(C)** Distribution of architectural patterns among transcriptomic regions in each patient; most patients display heterogeneous architectures. **(D)** Distribution of molecular clusters across spatial clusters (excluding patients with few regions), where each spatial cluster is transcriptionally homogeneous. **(E)** Distribution of architecture types across spatial clusters (excluding patients with few regions), with each cluster exhibiting consistent architectural profiles. **(F)** Spatial projections of tumor slides for patients 6 (top) and 3 (bottom), with color coding for architectures (left) and molecular clusters (right). **(G–I)** Heatmaps showing the top 50 differentially expressed genes across architecture types for crypt base (G), regenerative stem cell (H), and EMT (I) gene sets.

**Supplementary Fig. 3.**
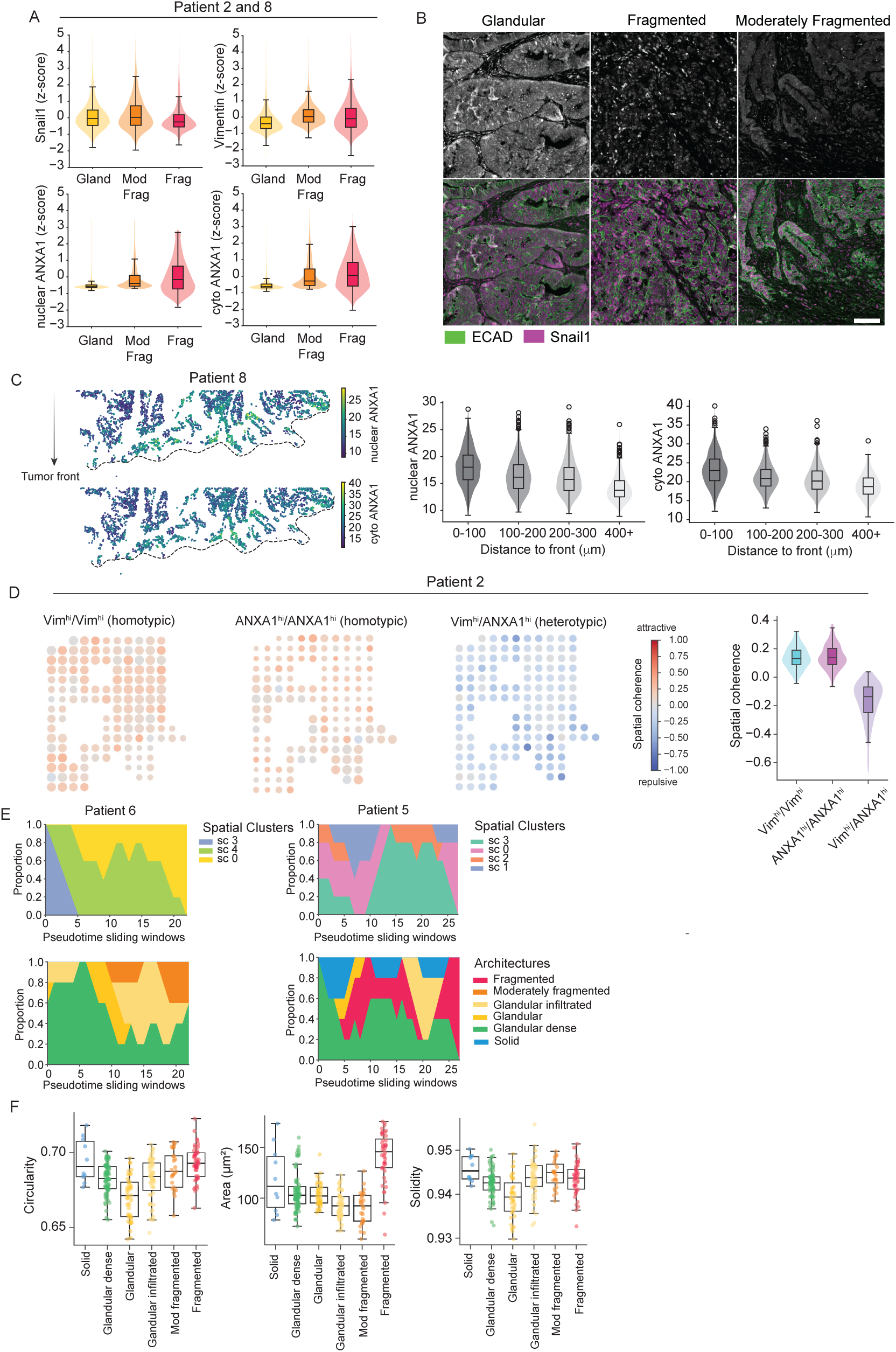
**(A)** Boxplots and violin plots showing IHC quantification of Snail1, Vimentin, and nuclear/cytoplasmic ANXA1 across three tumor architectures, derived from multiple regions of two whole-slide sections from two patients. Snail1 levels is expressed at similar across tumor architectures, whereas Vimentin and ANXA1 expression are elevated in fragmented regions. **(B)** Representative immunofluorescence images of E-cadherin (E-cad) and Snail1 stained tumor biopsies from glandular, moderately/highly fragmented regions. Scale bars 100 µm. **(C)** Projected mean intensity of nuclear (top) and cytoplasmic (bottom) ANXA1 in individual tumor cells at the invasive front of Patient 8 exhibiting moderate fragmentation (left). Corresponding boxplots and violin plots show ANXA1 intensity across distance bins relative to the invasive front (right). **(D)** Projected mean spatial-coherence score calculated for each tile, grouped by (i) homotypic pairs (cells of the same type) and (ii) heterotypic pairs (cells of different types). Positive scores reflect stronger-than-random co-localisation, whereas negative scores indicate spatial avoidance. In the homotypic pair plot, dot size scales with the relative abundance of each cell type. Boxplot and violin plots representing average spatial-coherence of each tile across different pairs. Note the clear segregation between the two populations: homotypic pairs cluster, while heterotypic pairs tend to remain apart. **(E)** Changes in spatial clusters (top) and architecture types (bottom) along pseudotime, profiled within consecutive sliding windows. (**F)** Boxplot representing nuclear shapes descriptors (Circularity, Area and Solidity) averaged per transcriptomic region.

**Supplementary Fig. 4.**
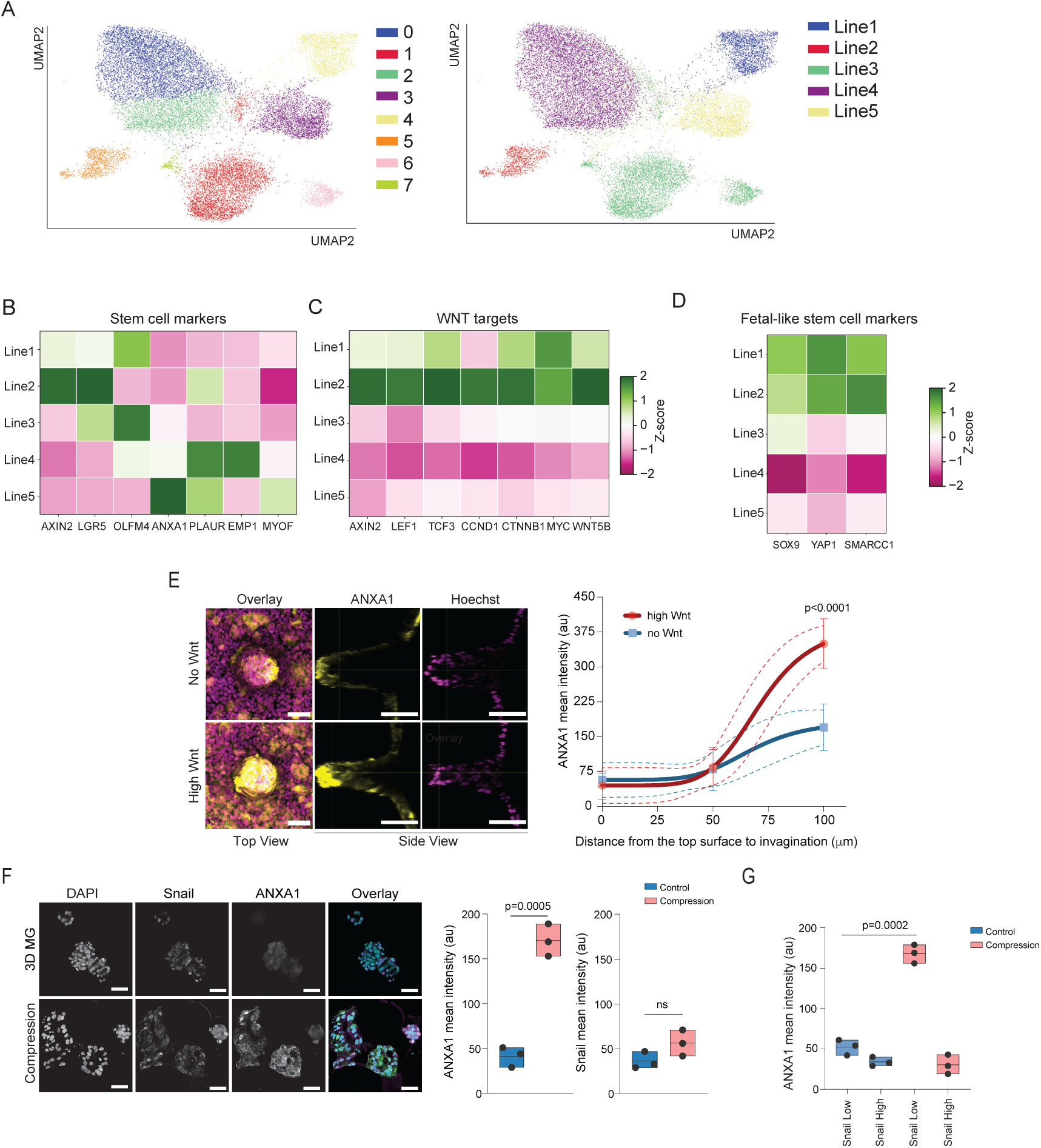
**(A)** UMAP projection of Leiden clusters from single cell RNAseq data of 5 distinct patient derived CRC organoids cultured in Matrigel. Right panel is color coded according to each line. **(B)** Heatmap of z-scores from mean expression levels of selected key stem cell marker genes across the cell lines. **(C)** Heatmap of z-scores from mean expression levels of selected key WNT target genes across the cell lines. **(D)** Heatmap of of z-scores from mean expression levels of selected fetal reprogramming factor genes across the cell lines. **(E)** Representative top- and side-view and quantification of ANXA1-stained patient-derived organoids from surface to invaginating architectures when grown in either high Wnt or no Wnt conditions. Scale bars 50 µm; n=3 independent experiments with 200 cells/condition/experiment from at least 3 structures; Two-Way Anova/Dunnet’s. **(F, G)** Representative images and quantification of ANXA1 and Snail intensity in mechanically compressed versus uncompressed organoids, as well as quantification of ANXA1 intensity binned by EMT status of the organoids as defined by binary levels of Snail (high or low) (G). Scale bars 30 µm; n=3 independent experiments with >300 cells/condition/experiment; Anova/Dunnet’s.

